# Heterogeneity of Long-History Migration Predicts Smiling, Laughter and Positive Emotion Across the Globe and Within the United States

**DOI:** 10.1101/317289

**Authors:** Paula M. Niedenthal, Magdalena Rychlowska, Adrienne Wood, Fangyun Zhao

**Affiliations:** Department of Psychology, University of Wisconsin-Madison, Madison, WI; School of Psychology, Queen’s University Belfast, Belfast, Ireland

## Abstract

Recent findings demonstrate that heterogeneity of long-history migration predicts present-day emotion behaviors and norms. Residents of countries characterized by high ancestral diversity display emotion expressions that are easier to decode by observers, endorse norms of higher emotion expressivity, and smile more in response to certain stimuli than residents of countries that lack ancestral diversity. We build on the extant findings and investigate historical heterogeneity as a predictor of daily smiling, laughter, and positive emotion across the world’s countries and the states of the United States. Study 1 finds that historical heterogeneity is positively associated with self-reports of smiling, laughter, and positive emotions in the Gallup World Poll when controlling for GDP and current present-day population diversity. Study 2 extends the findings to effects of long-history migration within the United States. We estimated the average percentage of foreign-born citizens in each state between 1850 and 2010 based on US Census information as an indicator of historical heterogeneity. Consistent with the world findings of Study 1, historical heterogeneity predicted smiling, laughter, and positive, but not negative, emotion. The relationships remained significant when controlling for per capita income and present-day diversity of population of each state. Together, the findings further demonstrate the important role of long-history migration in shaping emotion cultures of countries and states, which persist beyond the original socio-ecological conditions, and open promising avenues for cross-cultural research.

## Introduction

A trip to Indonesia or Nicaragua, an evening watching the Olympics on television, or a stroll through an American urban center each provides substantial evidence of great diversity in cultural practices across human societies. But just what are the origins of the cultural differences?

Recent accounts of culture, inspired by theories of biological evolution, propose one answer: Part of the global variations in human behavior and traits can be understood as selective adaptations to pressures posed by local social and ecological conditions (1,2). For example, evidence suggests that adjustment to the prevalence of pathogens in the immediate environment includes a reliance on authoritarian governing structures, the establishment of tight social norms especially about social interaction and sexuality, and individual-level traits of low extraversion and low openness to experience (3–5). These adaptations are part of the behavioral immune system (6) and become embodied as cultural practices and institutions serving to minimize exposure to diseases that are transmitted through inter-group contact.

The long-term demographic history of a population represents another socioecological condition to which systematic adaptations can be expected (7). Previous research links heterogeneity of long-term migration – a context associated with historical pressures to communicate in the absence of common language and social norms – with present-day emotion expressivity (8,9), the personality trait of openness to experience (10), and with the frequency of smiling in response to amusing or interesting stimuli (11). In the present work we build on these previous findings and use data from several global and national (within-U.S.) polling studies on emotional expressions and experiences to investigate how long-history migration patterns determine global and regional emotion cultures.

### Heterogeneity of Long History Migration

Beginning with migration out of Africa and bolstered by innovations that supported the rise of colonialization roughly 500 years ago, humans have dispersed across the globe in waves of often massive proportions (12). Some regions (e.g., present-day Argentina and New Zealand) received migrants from many, and others (e.g., Finland and South Korea) from far fewer, different cultural groups that were largely unknown to each other. Arguably, the pressures posed by the socio-ecological environment of cultural heterogeneity differ from those confronted in more homogeneous environments. In particular, survival in heterogenous societies relied on the exchange of unfamiliar concepts and practices, the formation and re-formation of social groups and hierarchies, and the creation of new institutions. According to a recent theory (7), the absence of norms for and shared language about abstract concepts such as emotion and motivation would have favored the reliance on nonverbal behavior in the service of social coordination.

Recent research provides evidence supporting the idea that heterogeneity of long history migration explains cross-cultural variations in emotion expression and experience. Wood and colleagues, for instance, investigated the relationship between historical heterogeneity and the transparency of people’s emotional expressions (9). The researchers re-analyzed existing findings from published studies that measured the accuracy of recognition of facial and vocal expressions of emotion across cultures (13). In each of the studies, spanning 92 articles involving participants from 79 cultures and expressions of representatives of 32 cultures, individuals from one culture were exposed to expressions of emotion of individuals from another culture. They then classified the expressions using a limited set of labels such as “joy” and “anger.”

Results revealed that the heterogeneity of the country of the expresser (but not the perceiver) was related to emotion recognition accuracy, such that expressers from historically heterogeneous cultures made displays that were easier to recognize across cultures. This finding supports the idea that a boost in the signal value of emotion in the face and the voice may constitute an adaptation to the pressure of interacting with individuals with whom one shares few expectations and no nuanced emotion language. In other words, living with people from diverse cultural backgrounds over time appears to be associated with the use of facial and vocal expressions that are relatively unambiguous and easily decoded by unfamiliar others.

Rychlowska and colleagues investigated the related prediction that, in contrast to members of homogeneous cultures, those in heterogeneous cultures would benefit from the cultural evolution of display rules that favor the spontaneous expression (versus dissimulation) of emotion (8). This reasoning follows from consideration of the social advantages of expressing one’s emotions (14), such as the establishment of trust through a transparency of communication and a facilitated understanding about the creation and achievement of goals. However, expressiveness is also costly and even risky, as it may disrupt social norms and existing hierarchies. Thus, display rules that favor emotional expressiveness make most sense in social contexts in which normative behavioral and emotional responses are not shared compared to the contexts in which expectations and cultural rules for emotion are widely shared (15).

To test this prediction, the authors reanalyzed an existing set of cross-cultural data about expressive display rules display rules governing the expression of anger, contempt, disgust, fear, happiness, sadness, and surprise across 32 countries (Matsumoto, Yoo, & Fontaine, 2008). Robust to the inclusion of other features of culture such as individualism and residential mobility in the statistical models, the findings revealed that display rules in heterogeneous cultures favor higher emotion expressivity than in homogeneous cultures (8).

Historical heterogeneity may also predict the expression of specific emotions. In particular, the smile should be more frequent in societies that have experienced heterogeneity of long-history migration, in order to signal lack of threat and establish trust in the service of successful cooperation (17–19). Initial evidence supporting this reasoning comes from a recent study in which 866,726 participants from 31 countries were filmed when watching video advertisements (11). The analysis of the recordings revealed that the proportion of video frames during which participants smiled was best predicted by historical heterogeneity. In other words, individuals from heterogeneous cultures spent significantly more time smiling than did those from homogeneous cultures. The effect was robust to the inclusion of other aspects of culture, such as urbanization, gross-domestic products, individualism (Hofstede, 2001), and ethnic fractionalization (Alesina et al., 2003).

### Overview of the Present Research

While the results of Girard and McDuff (2017) are consistent with the hypothesis that, in heterogeneous societies, certain types of smiles have been useful over cultural evolution in solving the social task of reinforcing behavior and inviting social interaction (8), they are not without limitations. Specifically, the study only examined smiles occurring in reaction to one type of stimuli (video advertisements) in a relatively restricted context of market research facilities. Thus, the findings may be due to the possibility that participants from homogeneous countries found the stimuli less entertaining or humorous than participants from heterogeneous countries. One purpose of the present research was to examine frequency of smiling in general. The Gallup World Poll and Gallup US Daily Tracking, which sampled individuals from a total of 150 nations and from 51 United States (including the District of Columbia), contained the question “Did you smile or laugh a lot yesterday?” While the item is memory-based, it allows for the possibility that smiling and laughter were elicited by any stimulus rather than only advertisements, especially that these behaviors are most common in social rather than solitary settings. We therefore analyzed this measure of smiling and laughter as a function of historical heterogeneity.

A second aim of the present research was to explore the implications of more frequent smiling and laughter, certain types of which are associated with positive emotions (20,21). Thus, frequent smiles and laughter could be an indication that members of heterogeneous cultures also experience more positive emotions than members of homogeneous cultures. To address this question, we examined the relationships between historical heterogeneity and measures of positive and negative emotions from several large-scale global and national surveys. Positive and negative emotional experiences are not strictly negatively correlated and have distinct relationships to features of culture and well-being (22). We examined negative emotions in order to be able to provide limiting conditions for the predictive value of historical heterogeneity.

Finally, the boundaries of existing nations do not inherently constitute the level at which the heterogeneity of cultures is defined. Regions within nations can vary in the extent to which they experienced extensive long history migration, a nuance that is lost in between-country analyses. Countries that are composed of provinces (Canada, Chile) or states (the United States) present opportunities to look at within-nation effects of heterogeneity on regional culture, since these nations maintain census data at the province/state level. In the present research we related the heterogeneity of the states of the United States to reports of smiles/laughter and positive experiences. Evidence that relationships hold at the level of the state would provide additional support for the significance of long term diversity of ancestry in establishing cultures of emotion (Nettle, 2009).

## Study 1

In Study 1, we built on the findings of Girard and McDuff (2017) and tested the hypothesis that country-level historical heterogeneity predicts smiling and laughter as well as positive and negative emotions. We used the number of source countries that contributed to the current population of a given country between 1500 and 2000 (Putterman & Weil, 2010) as an index of historical heterogeneity. The variable was then used as a predictor of smiling and laughter, four different measures of positive emotions, and five measures of negative emotions. We also included two control variables: gross domestic product (GDP) per capita and present-day migration diversity indexed by the number of countries contributing to the population of a given country in the year 2015. In the studies examining historical heterogeneity cited above (Rychlowska et al., 2015; Girard & McDuff, 2017; Shrira et al., 2018), this variable has been found to be a robust predictor of display rules for emotional expression, smiling, and also trait openness to experience when controlling for other aspects of culture (e.g., individualism) and economic conditions (e.g., GDP). In the present research we chose to control for GDP because of the complex relationship between economic viability and positive emotions (23). Measures of present-day diversity were included as controls in order to support conclusions about long-term versus more immediate effects of socioecological context on human behavior.

### Measures

#### Historical heterogeneity

Long-history heterogeneity vs. homogeneity of migration was operationalized by the number of countries in which the ancestors of a given country’s modern inhabitants lived in A.D. 1500. The index is derived from the World Migration Matrix (24), whose entries represent the fraction of a country’s ancestry in 1500 attributable to different source countries. The World Migration Matrix has been used in previous studies (9–11). The scores vary between 1 and 83 and are available for 172 countries. This variable will be called Heterogeneity and it is illustrated in Fig 1.

**Fig 1.**
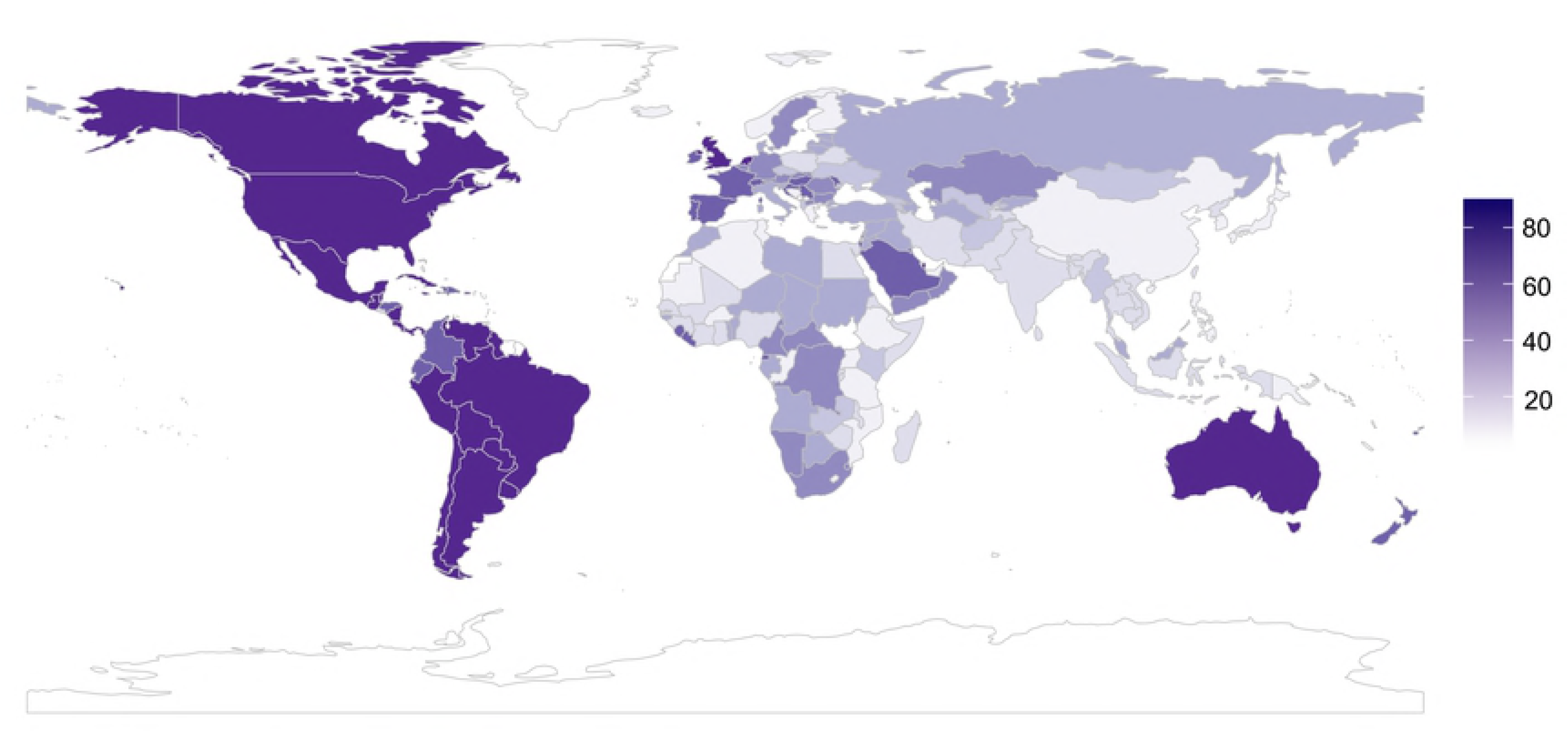
The number of source countries to the countries of the world since A.D. 1500. Darker colors indicate greater number of source countries. From Putterman and Weil (2010) World Migration Matrix.

#### Smiling and laughter

Measures of smiling and laughter were derived from the World Poll – a large international survey conducted by the Gallup Organization since 2005 in more than 160 countries, which samples over approximately 99% of the world’s adult population. Typically, at least 1000 respondents are polled in each country. The survey includes more than 100 global questions as well as region-specific items. Participants are interviewed by telephone or during face-to-face meetings. We used the latest available measures of smiling and laughter, from the 2017 Gallup World Poll (25) based on nearly 149,000 interviews with adults in 142 countries in 2016. The country-level scores reflect the percentage of respondents who answered “yes” to the question: “Did you smile or laugh a lot yesterday”. The measure was available for 142 countries, with scores ranging from 42 to 89%.

#### Measures of positive emotion

An Enjoyment measure was part of the 2017 Gallup World Poll. Similar to the index of smiling and laughter, the country-level scores reflect the percentage of respondents who answered “yes” to the question asking whether they experienced enjoyment during a lot of the day yesterday. Ratings were available for 142 countries and ranging from 33 to 91%.

A composite Positive Experience Index was computed for the purposes of the 2017 Global Emotions Report (26) using participants’ responses to five questions: “Did you feel well-rested yesterday?”, “Were you treated with respect all day yesterday?”, “Did you smile or laugh a lot yesterday?”, “Did you learn or do something interesting yesterday?” “Did you experience the following feelings during a lot of the day yesterday? How about enjoyment?”. The Positive Experience Index score is the mean of all valid affirmative responses to these items multiplied by 100. Scores were available for 142 countries and ranged from 0 to 100, with higher scores meaning that positive experiences are more pervasive in a country.

A measure of feelings of Happiness was derived from the World Values Survey (wave 6, 2010-2014; (27), a global research project covering 60 countries, with a minimum of 1000 respondents per country. The question “Taking all things together, would you say you are: 1) Very happy, 2) Rather happy, 3) Not very happy, 4) Not at all happy” was part of a large standardized questionnaire administered in face-to-face interviews and phone interviews for remote areas. Participants’ valid responses were averaged to obtain country-level measures. Scores were available for 60 countries and ranged between 1 and 4, with higher scores indicating lower levels of happiness.

Our measure of Positive Emotion, derived from the International College Survey 2001 (28) administered to 9857 college students in 48 countries, reflects the average frequency of positive emotions. In the survey, participants were asked to rate how often they had felt six positive emotions (pleasant, happy, cheerful, pride, gratitude, and love) using 9-point scales ranging from 1 (*not at all*) to 9 (*all the time*). Scores were averaged in a global measure of positive emotion, available for 46 countries, with higher values representing higher frequency of positive emotions.

#### Measures of negative emotion

We used measures of Anger, Sadness, Stress, and Worry, which were part of the 2017 Gallup World Poll. Country-level scores reflected the percentage of respondents who answered “yes” to the question asking whether they experienced anger, sadness, stress, and worry during a lot of the day yesterday. Ratings were available for 142 countries and ranged from 6 to 50% (anger), 7 to 61% (sadness), 12 to 66% (stress), and 15 to 74% (worry).

Similar to the Positive Experience Index described above, the Negative Experience Index was a composite measure of respondents’ well-being computed from five items asking whether participants experienced physical pain, worry, sadness, stress, and anger during a lot of the day yesterday. The Negative Experience Index score is the mean of all valid affirmative responses to these items multiplied by 100. The scores were available for 142 countries and ranged from 0 to 100, with higher scores indicating higher pervasiveness of negative experiences in a given country.

Similar to the Positive Emotion measure described above, the index of Negative Emotion was derived from the International College Survey 2001 (28) and averaged participants’ responses to the items assessing the frequency of eight negative emotions (sad, anger, unpleasant, guilt, shame, worry, stress, and jealousy). Scores were available for 46 countries, with higher values reflecting higher frequency of negative emotions.

#### GDP per capita

We used each country’s gross domestic product divided by its total population. Values for 2017 were retrieved from the World Factbook (29) and were available for 166 countries, ranging from $700 to $124,900.

#### Present-day migration diversity

The diversity of the present (vs. long-history) migration was indexed by the construct of ethnic fractionalization (30), reflecting the probability that two randomly selected individuals from a given country belong to different ethnic groups. Population data used to compute the variable were provided by the sources published between 1997 and 2001 or directly obtained from national censuses. Scores of ethnic fractionalization were available for 166 countries and varied between 0 and 1.

### Results

#### Correlations

We first calculated the relationships between historical heterogeneity, smiling and laughter, and measures of positive and negative emotion by computing pairwise correlations for all variables described earlier. An inspection of the correlation matrix (see Table 1) reveals significant positive associations between Heterogeneity and Smiling and Laughter, *r*(140) = .31, *p* < .001, 95% CI [.15, .45], Enjoyment, *r*(140) = .36, *p* < .001, 95% CI [.21, .50], Positive Experience Index, *r*(140) = .39, *p* < .001, 95% CI [.25, .52], and Positive Emotion, *r*(44) = .38, *p* < .01, 95% CI [.10, .60]. The correlation with the negatively coded measure of happiness was only marginally significant, *r*(58) = -.25, *p* = .05, 95% CI [-.47, .00]. Heterogeneity was significantly and negatively associated with the experience of Anger, *r*(140) = -.22, *p* < .01, 95% CI [-.37, -.06], but was not significantly related to other negative emotions, *r*s < .16, *p*s > .05. Additionally, there was a significant positive correlation between Heterogeneity and GDP per capita, *r*(164) = .25, *p* = .001, 95% CI [.10, .39].

**Table 1.**
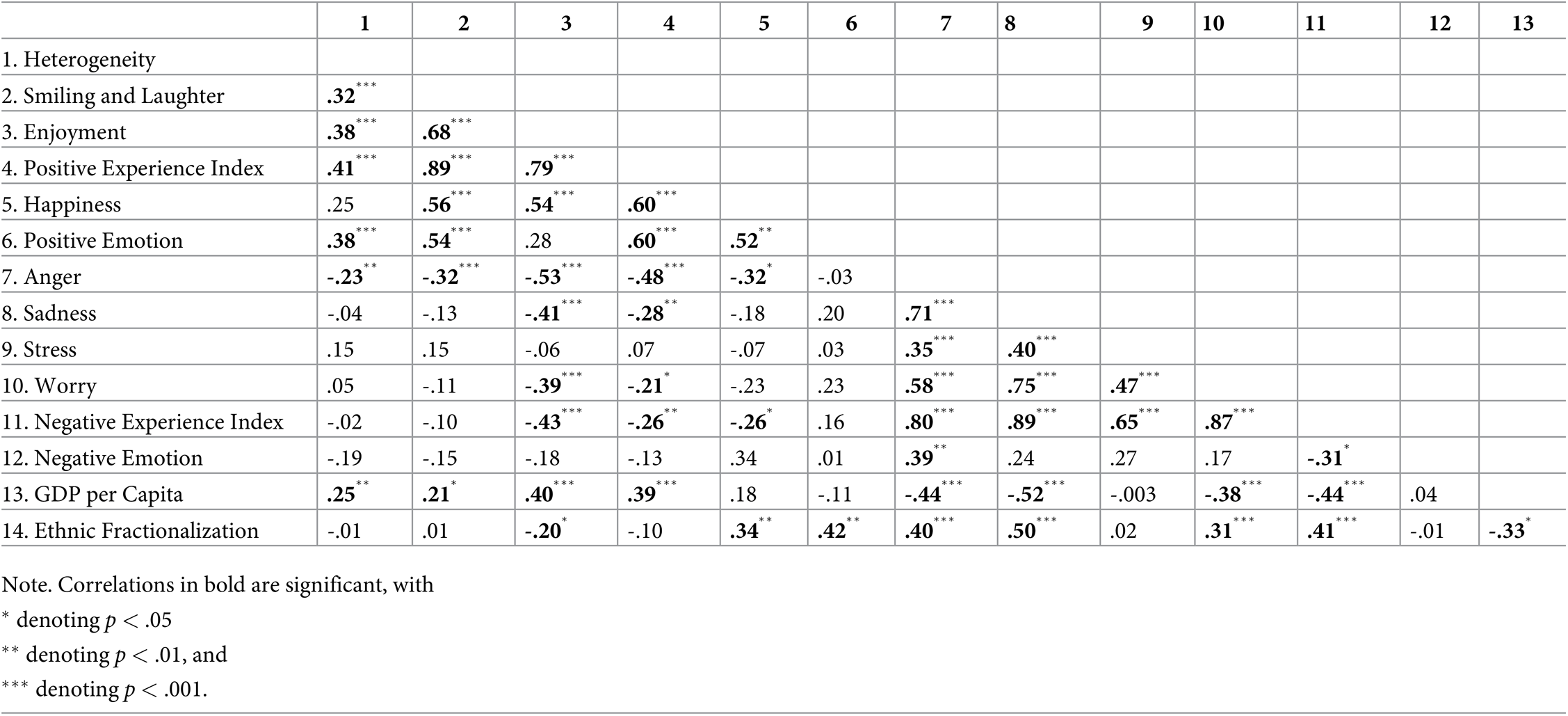
Correlations between variables in Study 1.

Overall, beyond predicting smiling and laughter, historical heterogeneity was also consistently correlated with measures of positive emotion. Correlations with negative emotions were less consistent, with only anger being significantly and negatively associated with Heterogeneity. To extend the correlation analyses, we conducted a series of multiple regressions examining the usefulness of Heterogeneity as a predictor of Smiling and Laughter, Enjoyment, and Positive Experience Index. The analyses focused on the measures derived from Gallup World Poll to maximize statistical power, as they included more data points than the World Values Survey and the International College Survey 2001. Each analysis included GDP per capita and Ethnic Fractionalization as control variables.

#### Predicting Smiling and Laughter

We first regressed the measure of Smiling and Laughter on GDP per capita and Ethnic Fractionalization, saving the standardized residuals. These residuals were then analyzed as a function of historical heterogeneity. In this analysis, Heterogeneity was a significant positive predictor of Smiling and Laughter, β = .28, *F*(1,139) = 11.43, *p* = .001, *R*^2^_*adj*_ = .07. The effect of Heterogeneity was also significant in a similar regression model, in which Ethnic Fractionalization was replaced by another index of present-day population diversity, namely the number of source countries contributing to the population of a given country in 2015 (31) (UN, 2018), β = .25, *F*(1,137) = 9.54, *p* = .002, *R^2^_adj_* = .06.

In a supplemental analysis, we simultaneously regressed the measure of Smiling and Laughter on Heterogeneity, GDP per capita, and Ethnic Fractionalization, *F*(3, 137) = 6.54, *p* < .001, *R^2^_adj_* = .11. Heterogeneity was a significant predictor of Smiling and Laughter, β = .28, *F*(1,137) = 11.77, *p* = .001. The same was true for GDP per capita: β = .19, *F*(1,137) = 4.31, *p* = .04. The effect of Ethnic Fractionalization was not significant, β = .10, *F*(1,137) = 1.17, *p* = .28.

#### Predicting Enjoyment

We first regressed the measure of Enjoyment on GDP per capita and Ethnic Fractionalization, saving the standardized residuals. A subsequent analysis with residuals as a dependent variable and Heterogeneity as a predictor revealed a significant effect of Heterogeneity on the residuals, β = .32, *F*(1,139) = 15.98, *p* < .001, *R^2^_adj_* = .10. The effect of Heterogeneity was also significant in the regression model in which Ethnic Fractionalization was replaced the number of source countries in 2015 (31), β = .30, *F*(1,137) = 13.22, *p* < .001, *R^2^_adj_* = .08.

A supplemental regression model including Enjoyment as a dependent variable and Heterogeneity, GDP per capita, and Ethnic Fractionalization as predictors, *F*(3, 137) = 15.20, *p* < .001, *R^2^_adj_* = .23, revealed a significant effect of Heterogeneity, β = .31, *F*(1,137) = 16.48, *p* < .001. The effect of GDP was also significant, β = .31, *F*(1,137) = 14.13, *p* < .001. Ethnic Fractionalization was not a significant predictor of Enjoyment, β = -.06, *F*(1,137) = 0.49, *p* = .48.

#### Predicting the Positive Experience Index

As in the previous analyses, Positive Experience Index was first regressed on GDP and Ethnic Fractionalization. A subsequent regression analysis revealed that the standardized residuals were significantly accounted for by Heterogeneity, *F*(1, 139) = 16.69, *p* < .001, *R^2^_adj_* = .12. The effect of Heterogeneity was also significant in the regression model in which Ethnic Fractionalization was replaced the number of source countries in 2015 (31), β = .30, *F*(1,137) = 13.22, *p* < .001, *R^2^_adj_* = .08.

A regression model including the Positive Experience Index as a dependent variable and Heterogeneity, GDP per capita, and Ethnic Fractionalization as predictors, *F*(3, 137) = 16.53, *p* < .001, *R^2^_adj_* = .25, revealed a significant effect of Heterogeneity, β = .34, *F*(1,137) = 20.31, *p* < .001. The effect of GDP per capita was also significant, β = .35, *F*(1,137) = 18.12, *p* < .001. Ethnic Fractionalization was not a significant predictor, β = .06, *F*(1,137) = 0.62, *p* = .43. Additional regression analyses, in which Heterogeneity and GDP per capita were log-transformed (using base 10) because of high kurtosis values yielded an identical pattern of results.

## Study 2

While Study 1 replicated and extended the findings of Girard and McDuff (2017) in showing that country-level historical heterogeneity is a positive predictor not only of smiling and laughter in a more general context, but also various indices of positive emotion, Study 2 focused on long-history migration within the United States. Historical heterogeneity of each of the U.S. states was estimated by averaging each decennial census’s percentage of foreign-born contributing to its population between 1850 and 2010. As in Study 1, we examined the significance of heterogeneity in predicting smiling and laughter, as well as six different measures of positive and negative emotions derived from the Gallup U.S. Daily Tracking Poll. We also included per capita income for each state and a measure of source countries contributing to each state’s population in 2016 as a measure of present-day diversity.

### Measures

#### Historical heterogeneity

Long-history heterogeneity of migration was operationalized as the percent of foreign-born citizens contributing to the population of each U.S. state between 1850 and 2010 provided by the US. Bureau of the Census. Scores were computed as an average of all census estimates available for a given state. Percentages of foreign-born individuals from 1850 to 2000 were retrieved from the decennial censuses (32,33) and from 2010 were part of the 2006-2010 American Community Survey (34), a questionnaire conducted by the U.S. Census Bureau that replaced the decennial censuses. Scores varied between 0.79 and 22.65% and were available for the 48 continental U.S. states as well as for Washington, D.C. Proportions of foreign-born citizens were available from 1860 for Kansas, North Dakota, Nebraska, Nevada, South Dakota, and Washington; from 1870 for Arizona, Colorado, Idaho, Montana, and Wyoming; and from 1890 for Oklahoma. We excluded Alaska and Hawaii from the analyses, as censuses from these states were only available starting in 1960, which would not provide the same long-history estimate as the other states. The variable will be called Heterogeneity and it is illustrated in Fig 2.

**Fig 2.**
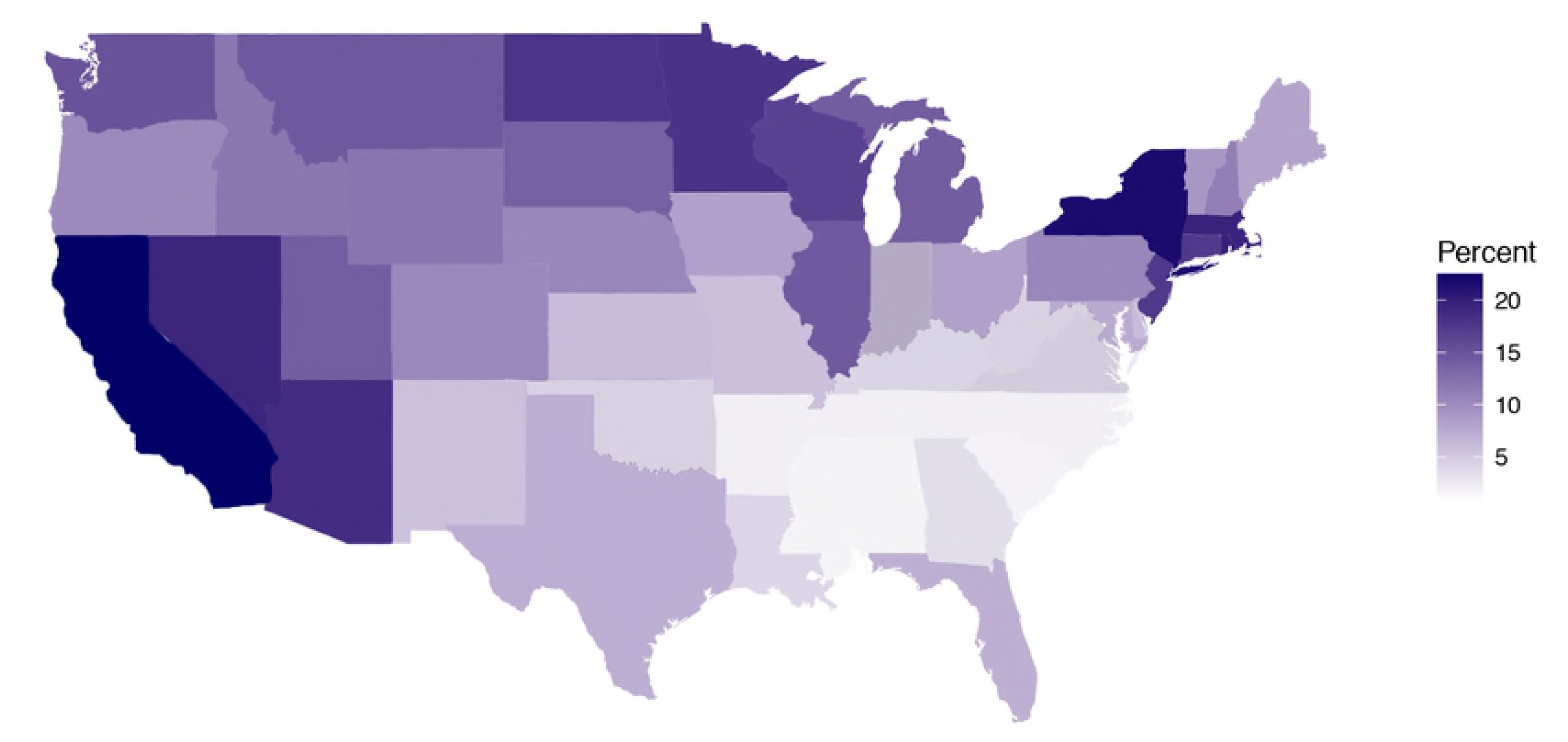
Average percent foreign born populations of the states of the continental United States. Based on censuses from 1850 (or between 1860 and 1890 for later-entry states) until 2010. Darker colors indicate higher percent foreign born populations.

#### Smiling and laughter

The measure of smiling and laughter was derived from the Gallup U.S. Daily Tracking survey, in which phone interviews are administered every day to approximately 500 randomly selected American respondents. The project yields large sample sizes, with as many as 175,000 participants surveyed each year. We used the latest available measures of smiling and laughter, from the 2016 U.S. Daily Tracking. The state-level scores reflected the percentage of respondents who answered “yes” to the question: “Did you smile or laugh a lot yesterday” and varied from 76 to 86%.

#### Measures of positive emotion

Similar to the index of smiling and laugher, measures of Enjoyment were also derived from the 2016 U.S. Daily Tracking Survey. The state-level scores indicated the percentage of respondents who answered “yes” to the question asking whether they experienced enjoyment during a lot of the day yesterday. Indications of enjoyment ranged from 82 to 91%.

The state-level Happoness scores indicated the percentage of respondents who answered “yes” to the question asking whether they experienced happiness during a lot of the day yesterday. Happiness scores ranged from 86 to 93%.

#### Measures of negative emotion

We analyzed reports of the emotions of anger, sadness, stress, and worry. The latest available measure of anger was collected as part of the 2013 U.S. Daily Tracking Survey and the three other measures were from 2016. State-level scores reflected the percentage of respondents who answered “yes” to the question asking whether they experienced anger, sadness, stress, and worry during a lot of the day yesterday. Endorsements ranged from 9 to 18% (anger), 11 to 20% (sadness), 36 to 46% (stress), and 23 to 36% (worry).

#### Income

We used state-level measures of income per capita. Values for 2016 were retrieved from the American Community Survey and ranged from $22,694 to $50,567.

#### Present-day migration diversity

The diversity of the present (vs. long-history) migration was indexed by the number of source countries whose citizens contributed to the population of a given state in the year 2016. Scores were retrieved from the report State Immigration Data Profiles (35) and ranged from 34 to 46.

### Results

#### Correlations

We first calculated the relationships between historical heterogeneity, smiling and laughter, and measures of positive and negative emotion by computing pairwise correlations between all variables. An inspection of the correlation matrix (see Table 2) reveals significant positive associations between Heterogeneity and Smiling and Laughter, *r*(47) = .46, *p* < .01, 95% CI [.20, .65] as well as Enjoyment, *r*(47) = .33, *p* = .02, 95% CI [-.06, .56], and Happiness, *r*(47) = .43, *p* < .01, 95% CI [.16, .63]. Heterogeneity was also negatively correlated with the measure of Sadness, *r*(47) = -.29, *p* = .04, 95% CI [-.53, -.01]. Additionally, there was a significant positive correlation between heterogeneity and income, *r*(47) = .47, *p* = .001, 95% CI [.22, .66]. None of the correlations between Heterogeneity and other negative emotions was significant, *r*s < .25, *p*s > .10.

**Table 2.** Correlations between variables in Study 2.

|  | 1 | 2 | 3 | 4 | 5 | 6 | 7 | 8 | 9 |
| --- | --- | --- | --- | --- | --- | --- | --- | --- | --- |
| 1. Heterogeneity |  |  |  |  |  |  |  |  |  |
| 2. Smiling and Laughter | **.46\*\*\*** |  |  |  |  |  |  |  |  |
| 3. Enjoyment | **.33\*** | **.69\*\*\*** |  |  |  |  |  |  |  |
| 4. Happiness | **.43\*\*\*** | **.70\*\*\*** | **.75\*\*\*** |  |  |  |  |  |  |
| 5. Anger | -.23 | **-.59\*\*\*** | **-.70\*\*\*** | **-.71\*\*\*** |  |  |  |  |  |
| 6. Sadness | **-.29\*** | **-.57\*\*\*** | **-.66\*\*\*** | **-.67\*\*\*** | **.63\*\*\*** |  |  |  |  |
| 7. Stress | .09 | -.19 | -.13 | -.07 | .23 | .33 |  |  |  |
| 8. Worry | -.01 | **-.35\*\*** | **-.41\*\*\*** | **-.36\*\*** | **.49\*\*** | **.65\*\*\*** | **.70\*\*\*** |  |  |
| 9. Income per capita | **.47\*\*\*** | **.32\*** | .16 | .09 | -.01 | -.17 | .17 | .23 |  |
| 10. Source countries in 2016 | .01 | -.10 | **-.35\*\*\*** | **-.44\*\*** | **.37\*\*** | **.46\*\*\*** | .09 | .19 | .16 |

Note. Correlations in bold are significant, with \* denoting $p < .05$ , \*\* denoting $p < .01$ , and \*\*\* denoting $p < .001$ .

We then conducted three multiple regressions examining the role of Heterogeneity as a spredictor of Smiling and Laughter, Happiness, and Enjoyment. Each analysis included state-level income and the number of source countries in 2016 as control variable.

#### Predicting Smiling and Laughter

We first regressed the measure of Smiling and Laughter on Income and the number of source countries in 2016, saving the standardized residuals. These residuals were then analyzed as a function of historical heterogeneity. In this analysis, Heterogeneity was a significant predictor of smiling and laughter, β = .32, *F*(1,47) = 5.45, *p* = .02, *R^2^_adj_* = .08. The effect of Heterogeneity was also significant in a similar regression model, in which Source countries in 2016 were replaced by another index of demographic diversity, namely the proportion of foreign-born citizens contributing to the population of a given state in 2015 (U.S. Census Bureau, 2011-2015), β = .30, *F*(1,47) = 4.81, *p* = .03, *R^2^_adj_* = .07.

A linear regression model including the measure of Smiling and Laughter as the dependent variable and Heterogeneity, Income, and the number of source countries in 2016 as predictors, *F*(3, 45) = 4.77, *p* = .01, *R^2^_adj_* = .19, revealed a significant effect of Heterogeneity, β = .39, *F*(1,45) = 6.97, *p* = .01. The effects of Income and Source countries in 2016 were not significant, β = .15, *F*(1,45) = 1.08, *p* = .30 and β = -.13, *F*(1,45) = 0.98, *p* = .33, respectively.

#### Predicting Enjoyment

Again, we regressed the measure of Enjoyment on Income and the Source countries in 2016, saving the standardized residuals. These residuals were then regressed on historical heterogeneity. Heterogeneity was a marginally significant predictor of Enjoyment, β = .26, *F*(1,47) = 3.34, *p* = .07, *R^2^_adj_* = .05. The effect of Heterogeneity was also significant in the regression model, in which Source countries in 2016 were replaced by the proportion of foreign-born citizens in 2015, β = .36, *F*(1,47) = 6.89, *p* = .01, *R^2^_adj_* = .11.

A linear regression model including the measure of Enjoyment as the dependent variable and Heterogeneity, Income, and the number of source countries in 2016 as predictors, *F*(3, 45) = 4.74, *p* = .01, *R^2^_adj_* = .19, revealed a significant effect of Heterogeneity, β = .30, *F*(1,45) = 4.21, *p* = .05. The number of source countries in 2016 was negatively associated with Enjoyment, β = -.36, *F*(1,45) = 7.58, *p* = .01, The effect of Income was not significant, β = .08, *F*(1,45) = 0.29, *p* = .60.

#### Predicting Happiness

Again, we regressed the measure of Happiness on Income and the Source countries in 2016, saving the standardized residuals. These residuals were then regressed on historical heterogeneity. Heterogeneity was a significant predictor of Happiness, β = .40, *F*(1,47) = 9.04, *p* < . 01, *R^2^_adj_* = .14. The effect of Heterogeneity was also significant in the regression model, in which Source countries in 2016 were replaced by the proportion of foreign-born citizens in 2015 (U.S. Census Bureau, 2011-2015), β = .44, *F*(1,47) = 11.51, *p* = .001, *R^2^_adj_* = .18.

A linear regression model including the measure of Happiness as the dependent variable and Heterogeneity, Income, and the number of source countries in 2016 as predictors, *F*(3, 45) = 9.24, *p* < .001, *R^2^_adj_* = .34, revealed a significant effect of Heterogeneity, β = .46, *F*(1,45) = 11.82, *p* = .001. The number of source countries in 2016 was negatively associated with Happiness, β = -.44, *F*(1,45) = 13.40, *p* = .001, The effect of Income was not significant, β = -.05, *F*(1,45) = 0.15, *p* = .70.

## General Discussion

In two studies, we tested predictions derived from the idea that long-history migratory patterns resulting in high ancestral heterogeneity constitute a force that determines long-lasting aspects of emotion culture. The need to coordinate and build societal institutions in the absence of shared initial language and emotion norms creates a context with strong pressures for efficient non-verbal communication. Recent studies have examined behaviors such as smiling while viewing advertisements (11) and facial expression recognition accuracy (9) and have found support for the significance of this socioecological factor: Individuals from historically heterogeneous cultures smile more and display facial expressions that are more readily decoded across cultures. The analysis of the large-sample datasets reported in the present work is compelling because the approach allows us to consider most of the world’s countries rather than a subset and to explore additional emotion experiences. We also extended previous findings by examining indicators of smiling and positive emotions not bound to specific contexts or stimuli.

Using responses to the Gallup World Poll query about smiling and laughing on the previous day, we replicated the finding that historical heterogeneity is related to the frequency of smiling (11). Robust to the inclusion of GDP and present-day population diversity, historical heterogeneity was positively associated with reports of smiling and laughter on the previous day. An extant cross-cultural study found that signaling non-treat and openness to affiliation was a more important determinant of smiling for members of heterogeneous cultures than for members of homogeneous cultures (Rychlowska et al., 2015). It is thus possible that the more frequent smiling in historically heterogeneous countries observed in the present study reflects an adaptation to societal pressures to use nonverbal behavior – such as smiling – to invite and reinforce social interaction and cooperation.

Historical heterogeneity also predicted enjoyment and positive experiences more generally: Members of heterogeneous countries reported that they had felt enjoyment and had positive experiences on the previous day with greater frequency than members of homogeneous countries. These findings, consistent with those for smiles and laughter, were found over and above any effects of GDP and present-day population diversity. While present-day diversity was a significant predictor in some analyses, its effects were not as consistent as the effects of historical heterogeneity. Specifically, the effects of Ethnic Fractionalization in the analyses of the world data were not significant. However, the number of source countries in 2016, used as a control variable in the within-US analyses, was a significant predictor of enjoyment and happiness. Importantly, while historical heterogeneity was positively associated with the measures of positive emotions, present-day population diversity was negatively associated with these measures suggesting that the two socioecological variables have different effects. While long-history migration may encourage specific emotion behaviors and reactions and shape an emotion culture over centuries (Cohen, 2001), present-day population movements may represent initial conditions, which exert their immediate social and economic effects, but are not yet incorporated in societal institutions and norms.

Because of the correlational nature of the present findings, we cannot draw strong conclusions about how or why historical heterogeneity is related to the experience of positive emotion and experience. As suggested, these outcomes could be related to the frequent use of smiles. A reliance on smiling to invite and maintain channels for new relationships can have salutary effects on emotions. This is because facial expressions can feed back to modulate emotional experiences (36,37). As an example, when participants in one study were covertly induced to smile while undergoing a painful cold-pressor task, they showed lower physiological arousal and self-reported negative affect than control participants who completed the task with a neutral expression (38). In addition to these beneficial intrapersonal effects of smiles, observers make positive attributions about smiling people and trust them more than people who do not express smiles (19). Smiles thus provide one basis for successful cooperation (17). As a consequence, positive experiences may result from a cultural adaptation that involves greater use of the smile.

There is no formal reason to estimate historical heterogeneity only at the level of the country. Putterman and Weil’s World Migration Matrix (2010) provides such information, and the indicators of the number of source countries to present-day populations derived from the matrix has now been used in numerous studies. However, heterogeneity can also be analyzed within smaller territories of the same country. In the present research, we estimated the historical heterogeneity of the states of the United States because the states constitute territories that differ in their long-history migration. Our indicator of historical heterogeneity was an average of the percentage of foreign born population from the US Census from 1850 (or beginning when the state entered the Union) until 2010. This 160-year period is not as extended as the 500-year one that can be estimated for the world’s countries but does reflect an important portion of the migratory history of the United States during which numerous large-scale waves of immigration from diverse countries contributed to the overall populations of many states. The natural waterways and the agricultural opportunities offered by the geographic conditions resulted in high variability in these percentages.

Using the Gallup US Daily Tracking surveys, we were able to perform analyses similar to those that we conducted for the world as a whole. Our results were overall consistent with the findings from the analyses of the countries of the world. Historical heterogeneity of the states of the Unites States was predictive of smiling and laughter such that, again, residents of more historically heterogeneous states reported more smiles and laugher. In addition, the positive relationship extended to reports of enjoyment and happiness. Overall, historical heterogeneity was related to higher levels of positive emotion. In the US sample, present-day diversity was also significantly and negatively related to positive experiences of happiness and enjoyment.

Future work should investigate potential causes of the relationship between historical heterogeneity and the experience of positive emotions. Ongoing experimental work in our lab is examining how specific socio-ecological factors associated with heterogeneity – for instance a lack of shared verbal language boosting people’s reliance on nonverbal communication – lead to shifts in expressive behavior. Such is the strength of a socioecological perspective on cross-cultural differences: if specific features of the social environment exert the hypothesized influence on behavior, it should be observable in the laboratory and in prospective studies of cultural change.

## Acknowledgements

P.M.N. was supported by a 2017-2018 Cattell Sabbatical Award (MSN206694) from the James McKeen Cattell Fund. A.W. was support by an Emotion Research Training Grant (T32MH018931-24) from the National Institute of Mental Health. We thank Markus Brauer for discussions of and advice on correct data-analytic strategies and Elizabeth Edwards and Dafydd Roberts for their help with retrieving census data and the Nuffield Foundation for funding their work.

## References

1. Oishi S. Socioecological Psychology. Annu Rev Psychol. 2014 Jan 3;65(1):581–609.

2. Sng O, Neuberg SL, Varnum MEW, Kenrick DT. The Behavioral Ecology of Cultural Psychological Variation. Psychol Rev. 2018;

3. Gelfand MJ, Raver JL, Nishii L, Leslie LM, Lun J, Lim BC, et al. Differences Between Tight and Loose Cultures: A 33-Nation Study. Science. 2011 May 26;332:1100–4.

4. Murray DR, Schaller M. Historical Prevalence of Infectious Diseases Within 230 Geopolitical Regions: A Tool for Investigating Origins of Culture. J Cross-Cult Psychol. 2010 Jan;41(1):99–108.

5. Murray DR, Schaller M, Suedfeld P. Pathogens and Politics: Further Evidence That Parasite Prevalence Predicts Authoritarianism. PLOS ONE. 2013 May 1;8(5):e62275.

6. Schaller M, Park JH. The Behavioral Immune System (and Why It Matters). Curr Dir Psychol Sci. 2011 Apr;20(2):99–103.

7. Niedenthal PM, Rychlowska M, Wood A. Feelings and contexts: socioecological influences on the nonverbal expression of emotion. Curr Opin Psychol. 2017 Oct 1;17:170–5.

8. Rychlowska M, Miyamoto Y, Matsumoto D, Hess U, Gilboa-Schechtman E, Kamble S, et al. Heterogeneity of long-history migration explains cultural differences in reports of emotional expressivity and the functions of smiles. Proc Natl Acad Sci. 2015;112(19):E2429–36.

9. Wood A, Rychlowska M, Niedenthal PM. Heterogeneity of long-history migration predicts emotion recognition accuracy. Emotion. 2016;16:413–20.

10. Shrira I, Wisman A, Noguchi K. Diversity of historical ancestry and personality traits across 56 cultures. Personal Individ Differ. 2018 Jul;128:44–8.

11. Girard JM, McDuff D. Historical Heterogeneity Predicts Smiling: Evidence from Large-Scale Observational Analyses. In IEEE; 2017 [cited 2018 Apr 15]. p. 719–26. Available from: http://ieeexplore.ieee.org/document/7961812/

12. Diamond J. Guns, germs, and steel: the fates of human societies. NY: WW Norton & Company 14; 1997.

13. Elfenbein HA, Ambady N. On the universality and cultural specificity of emotion recognition: A meta-analysis. Psychol Bull. 2002;128:203–35.

14. Schug J, Matsumoto D, Horita Y, Yamagishi T, Bonnet K. Emotional expressivity as a signal of cooperation. Evol Hum Behav. 2010 Mar;31(2):87–94.

15. Matsumoto D, Yoo SH, Chung J. The Expression of Anger Across Cultures. In: Potegal M, Stemmler G, Spielberger C, editors. International Handbook of Anger [Internet]. New York, NY: Springer New York; 2010 [cited 2018 Apr 20]. p. 125–37. Available from: http://link.springer.com/10.1007/978-0-387-89676-2_8

16. Matsumoto D, Yoo SH, Fontaine J. Mapping Expressive Differences Around the World: The Relationship Between Emotional Display Rules and Individualism Versus Collectivism. J Cross-Cult Psychol. 2008 Jan;39(1):55–74.

17. Manzini P, Sadrieh A, Vriend NJ. On Smiles, Winks and Handshakes as Coordination Devices. Econ J. 2009;119(537):826–54.

18. Mehu M, Little AC, Dunbar RIM. Duchenne smiles and the perception of generosity and sociability in faces. J Evol Psychol. 2007 Mar;5(1):183–96.

19. Todorov A, Pakrashi M, Oosterhof NN. Evaluating Faces on Trustworthiness After Minimal Time Exposure. Soc Cogn. 2009 Dec;27(6):813–33.

20. Martin J, Rychlowska M, Wood A, Niedenthal P. Smiles as Multipurpose Social Signals. Trends Cogn Sci. 2017 Nov;21(11):864–77.

21. Wood A, Martin J, Niedenthal P. Towards a social functional account of laughter: Acoustic features convey reward, affiliation, and dominance. PLOS ONE. 2017 Aug 29;12(8):e0183811.

22. Kuppens, P, Realo A, Diener, E.The role of positive and negative emotions in life-satisfaction judgment across nations. J Pers Soc Psychol. 2008;66–75.

23. Easterlin RA, McVey LA, Switek M, Sawangfa O, Zweig JS. The happiness–income paradox revisited. Proc Natl Acad Sci. 2010 Dec 28;107(52):22463–8.

24. Putterman L, Weil DN. Post-1500 population flows and the long-run determinants of economic growth and inequality. Q J Econ. 2010;125(4):1627–82.

25. Gallup Analytics. (2018). Available from: http://www.gallup.com/analytics/213617/gallup-analytics.aspx.

26. Gallup. (2018). Gallup 2017 Global Emotions. Retrieved from:http://news.gallup.com/reports/212648/gallup-global-emotions-report-2017.aspx

27. Inglehart, R., C., Haerpfer A., Moreno, C., Welzel, K., Kizilova, J., Diez-Medrano, M., Lagos, P., Norris, E., Ponarin & B., Puranen et al. (eds.). 2014. World Values Survey: Round Six-Country-Pooled Datafile Version:www.worldvaluessurvey.org/WVSDocumentationWV6.jsp. Madrid: JD Systems Institute.

28. Kuppens P, Ceulemans E, Timmerman ME, Diener E, Kim-Prieto C. Universal intracultural and intercultural dimensions of the recalled frequency of emotional experience. J Cross-Cult Psychol. 2006 Sep;37(5):491–515.

29. The World Factbook — Central Intelligence Agency [Internet]. [cited 2018 May 3]. Available from: https://www.cia.gov/library/publications/the-world-factbook/rankorder/2004rank.html

30. Alesina A, Devleeschauwer A, Easterly W, Kurlat S, Wacziarg R. Fractionalization. J Econ Growth. 8(2):40.

31. United Nations Population Division | Department of Economic and Social Affairs Available from:http://www.un.org/en/development/desa/population/migration/data/estimates2/estimates15.shtml

32. U.S. Bureau of the Census. (1999). Nativity of the population, for regions, divisions, and states: 1850 to 1990. Retrieved from: https://www.census.gov/population/www/documentation/twps0029/tab13.html

33. United States Census Bureau. (2003). The foreign-born population: 2000. Census 2000 Brief. Retrieved from: https://www.census.gov/prod/2003pubs/c2kbr-34.pdf

34. United Status Census Bureau. (2006-2010). Percent of People who are foreign born, 2006-2010 American Community Survey 5-year estimates. Retrieved from: https://factfinder.census.gov/faces/tableservices/jsf/pages/productview.xhtml?pid=ACS_10_5YR_GCT0501.ST04&prodType=table

35. Migration Policy Institute. (2018). State Immigration Data Profiles. Originally published on the Migration Policy Institute’s Migration Data Hub. (www.migrationpolicy.org/programs/migration-data-hub)

36. Price TF, Peterson CK, Harmon-Jones E. The emotive neuroscience of embodiment. Motiv Emot. 2012 Mar;36(1):27–37.

37. Wood A, Rychlowska M, Korb S, Niedenthal P. Fashioning the Face: Sensorimotor Simulation Contributes to Facial Expression Recognition. Trends Cogn Sci. 2016 Mar;20(3):227–40.

38. Kraft TL, Pressman SD. Grin and Bear It: The Influence of Manipulated Facial Expression on the Stress Response. Psychol Sci. 2012 Nov;23(11):1372–8.

